# Drone-Based Identification of Flood-Tolerant Maize via Multispectral Imaging: A Real-World Case Study

**DOI:** 10.1101/2024.11.26.625493

**Authors:** Madison Mitchell, Grace Sidberry, Morgan Mathison, Aaron J. DeSalvio, Daniel Kick, Jacob D. Washburn

## Abstract

Excess moisture (flooding, water logging, etc.) is a major source of crop damage causing catastrophic monetary losses to farmers around the world. Losses from excess water are often more common and costly than those from too little water (i.e., drought). Extreme weather patterns are predicted to increase, increasing the expected frequency of excess moisture events to farmers across the Midwest. Despite its importance, studying the impacts of flooding in the field is challenging due to unpredictability of flooding and fields being rendered inaccessible during flooding. Here, we took advantage of a natural flood experiment to examine the responses, damage, and recovery of diverse maize hybrids. Using drones, we monitored the hybrids before, during, and after flooding and examined the spatial and genetic components associated with post-flood survival.

## 1 INTRODUCTION

Excess water (e.g., floods, rain, etc.) was responsible for more indemnity payments than drought in midwestern states in 6 out of 10 years from 2014-2023 (United States Department of Agriculture - Risk Management Agency n.d.). Accurately assessing these damages and understanding the genetic basis for survival under these conditions is challenging, especially in the field for flood damage where the fields may become inaccessible both during and after the flooding event. Excess moisture events are also predicted to increase as climate change brings more extreme weather (Rosenzweig et al. 2002; Southworth et al. 2000). Unoccupied Aerial Vehicles (UAVs or Drones) are becoming a common phenotyping method because they remove these challenges, and in our case address safety risks relative to manually measuring flooded crops, all while maintaining or increasing image resolution and accuracy (Sweet et al. 2022) as compared to satellite data.

Numerous associations have been documented between waterlogging and flood stress to root characteristics (Jiménez et al. 2024; Mano et al. 2006; Ren et al. 2016). These include the presence of aerenchyma, stunted root growth, and decay (Kaur et al. 2019; Mano et al. 2006). Characteristics of flooded maize (*Zea mays*) at various growth stages have shown that nutrient absorption decreases under waterlogging and causes leaf area and photosynthetic capacity to decrease (Kaur et al. 2019; Ren et al. 2016). In most cases, excessive moisture limits oxygen availability to the roots, reducing root mass and increasing root death (Victor McDaniel, R. Wayne Skaggs, and Lamyaa M. Negm 2016). In a study looking at excessive moisture effects on roots, controlled flood conditions for five days led to maize root mortality rates ranging from 43 to 100% across all observed growth stages (Victor McDaniel et al. 2016). Yearly flooding lowers overall crop yields, costing farmers billions of dollars (>170) annually (Razzaq et al. 2021).

In addition, flooding and the effects of waterlogged soils and heavy silt deposits left on plants after a flood can result in lodging. In maize, lodging can occur in the form of stalk lodging, where breakage is between the ground and the top ear, root lodging, where plants lean more than 15% from completely vertical, or both (Genomes to Fields 2024). Root lodging occurs most often as a result of early-season storms that occur before maize plants develop brace roots (Tirado, Hirsch, and Springer 2021). The plants often fall over as a result of roots being partially pulled out of the soil (Lindsey, Carter, and Thomison 2021). While partial recovery is possible through “goose necking,” where plants bend towards the sun after being previously lodged (Genomes to Fields 2024; Tirado et al. 2021), root lodging reduces the photosynthetic productivity of the plant, limiting yield (Tirado et al. 2021). Yield losses in root-lodged fields can result in yield reductions of up to 28% in the V12 growth stage (Liu et al. 2023). L.-x. Tian, et. al. found that flooding of maize plants at the V3 stage negatively impacted the vascular bundle structure of the stem, which decreased the plants’ lodging resistance (Tian et al. 2020). Even though various growth stages have been analyzed, few have included data from each week of the growing season or found a high-throughput option for outdoor phenotyping.

Moderate Resolution Imaging Spectroradiometer (MODIS), an imaging sensor attached to a satellite orbiting Earth, has been deployed to gather information at the county level and compare flood and non-flood years (Shrestha et al. 2013). The authors found that flood years showed decreased normalized difference vegetative index (NDVI) values between late vs. early season flooding on a county level using 12 years of data (Shrestha et al. 2013). They also showed the possibility of recovery from replanting fields (Shrestha et al. 2013). The use of drones, rather than satellites, offers a higher-resolution image of the field allowing one to look at individual plots for specific characteristics and assess genetic differences (Sweet et al. 2022). Many useful manual measurements of phenotype can be collected on the ground, including plant height, ear height, days to maturity, and others (Genomes to Fields 2024). However, these take significant amounts of time and labor as compared to UAVs and potentially introduce experimenter effects, not to mention introducing safety risks in flooded conditions.

Here we describe the utility of both standard red, green, blue (RGB), and multi-spectral UAV imaging from before and after an unexpected natural flooding event for characterizing the influences of both the plot location within a field and genetic factors associated with flood survival and regrowth in maize. We developed a pipeline for extracting useful phenotypes from pre- and post-flood fields. This included the creation of orthomosaic images and extraction of traits like the Normalized Difference Vegetation Index (NDVI) and lodging from periods spanning the entire growing season. A comprehensive view of each genotype’s performance was visualized and statistically analyzed to assess whether genotypes were successful. Comparisons were made between the flooded field and the same genetic material in an unflooded environment. We found both spatial and genetic influences on survival and regrowth, as well as a significant and intriguing GWAS hit near genes involved in plant hormone regulation. This information provides a window into the responses of diverse maize genetics to flood scenarios and insight into how one might breed for more flood-tolerant crops in the future.

## 2 METHODS AND MATERIALS

### 2.1 Planting

This study used two field experiments with the same genetic materials in different years and different locations. One experiment, conducted at the Rollins Farm in Columbia, MO in 2021 experienced a natural flooding event. The other experiment, conducted at the Bradford Research Farm in Columbia, MO in 2020 did not experience flooding and was used as a comparison for the analyses. Both experiments contained the same 410 maize hybrids and were originally designed and conducted in collaboration with the Genomes to Fields (G2F) initiative Genotype by Environment (GxE) project.

The characteristics of the Rollins Farm Field location included Haymond Silt Loam, a commonly flooded soil order (United States Department of Agriculture - Natural Resources Conservation Service n.d.). Both fields were planted in a modified randomized complete block design with 20ft two-row plots and hybrid guard rows planted around the field. Within plots, 70 seeds were planted (35 per row) at a depth of 2 inches using a Winterstiger Plot King vacuum planter.

### 2.2 Flooding Scheme

In 2021 Rollins Farm research fields (Fig. 1A) were entirely submerged by the highest water levels on record from the nearby Hinkson Creek. The flooding lasted for approximately 24 hours. Before the flood, the maize within the experiment was at approximately the V5 maturity stage (Fig. 1B). Water flowed from Hinkson Creek to a portion of low-lying land and then over the levee into the research plots (Fig. 1A). Once the water had breached the levee it completely covered the plants to a height of several feet above them (Fig. 1C). After 24 hours, the majority of standing water had receded, but the soil remained waterlogged for at least a week longer.

**Figure 1:**
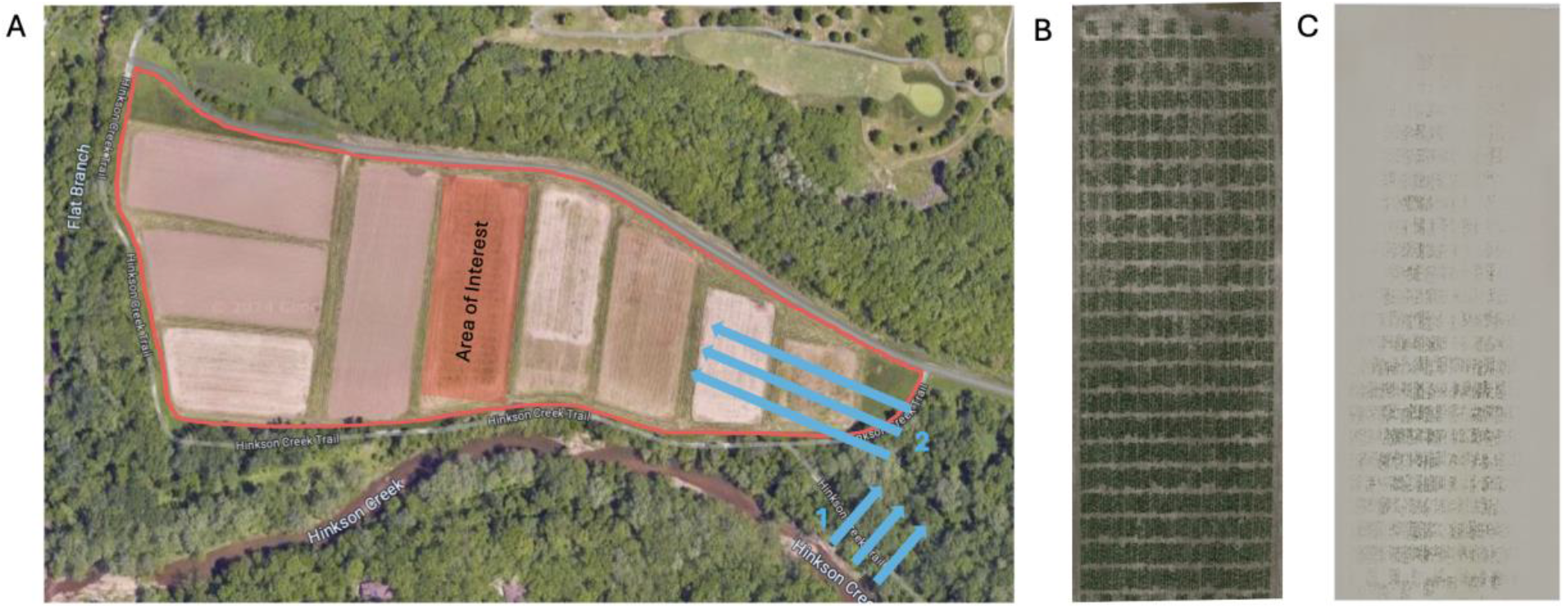
A) Map of the water infiltration in the planting zone of the 2021 flooded field, the red line denotes the dike that surrounds the farm, blue arrow 1 represents the first movement of water into a low-lying portion of land, blue arrow 2 shows the direction of water infiltration into the farmland once water levels rose above the height of the dike. B) the field before the flooding event occurred C) the completely submerged field during the peak of the flood

### 2.2 Data Collection

#### 2.2.1 Drone Imaging

For the whole growing season, weekly RGB and multispectral images were taken via UAV. RGB images were captured using a Mavic 2 Pro (Da-Jiang Innovations). NDVI images were collected throughout the season using a Matrice 600 Pro (Da-Jiang Innovations) UAV and RedEdge-MX (Micasense) multispectral camera. Images were collected between 11:00 am - 3:00 pm CST (around solar noon) to reduce the amount of shadow created by plants and increase the accuracy and uniformity of the data.

#### 2.2.2 Lodging Measurement

G2F Standard Operation Procedures: https://www.genomes2fields.org/resources/

### 2.3 Orthomosaic Creation

#### 2.3.1 Agisoft

RGB orthomosaics were processed in Agisoft Metashape Version 2.0.0 (Agisoft LLC, St. Petersburg, Russia). A summary of the workflow is as follows: 1) RGB images from the Mavic 2 Pro were loaded into Metashape; 2) photo alignment was conducted using the referenced preselection option with a key point limit of 40,000 and a tie point limit of 4,000; 3) initial bundle adjustment was conducted using the f, cx/cy, k1, k2, k3, p1, and p2 distortion parameters; 4) ground control points (GCPs) were imported into the Metashape project from a CSV file and their locations were tagged in six images per GCP; 5) in the reference pane, the images were deselected and the GCPs were integrated into the sparse point cloud (referred to as “tie points” in Metashape versions > 2.0.0) using the “update” button; 6) the dense point cloud (simply referred to as the “point cloud” in Metashape versions > 2.0.0) and digital elevation map (DEM) were built using the default parameters; 7) the orthomosaic was generated with the DEM as the surface.

Multispectral orthomosaics with five bands (blue, green, red, red edge, and near infrared) were also created in Metashape according to a protocol generated by the Agisoft Helpdesk Portal in conjunction with MicaSense recommendations (Agisoft Helpdesk Portal, 2024). The MicaSense RedEdge-MX camera saves five TIFFs each time a picture is taken, requiring raw images in Metashape to be imported using the “multi-camera system” option, which automatically reads the metadata of each image and ensures that all five bands are associated with a single image. After importing the images, reflectance calibration panel images are sorted into a separate folder automatically. The remaining steps are as follows: 1) reflectance calibration was conducted in the Tools > Calibrate Reflectance window using a CSV file with calibration information obtained from MicaSense that is specific to the panel that was purchased. Both the reflectance panels and sun sensor options enabled during reflectance calibration; 2) photo alignment and camera optimization were identical to the RGB workflow except generic preselection was used instead of referenced; 3) the point cloud was built with aggressive depth filtering; 4) the DEM and orthomosaic were built using default parameters; 5) to obtain reflectance values, all five bands were divided by 32768 (which is equal to 100% reflectance) by navigating to Tools > Set Raster Transform. Here, each output band labeled (B1-B5) was divided by 32768 with the “Enable transform” option selected; 6) the orthomosaic was exported with the “Raster transform: index value” option enabled to ensure the normalized values were saved.

#### 2.3.2 QGIS

QGIS was used to create bounding boxes around the individual orthomosaic images and save them for later use in data extraction within R (QGIS.org). The QGIS protocol used can be found at https://mnmzbd24.github.io/posts/QGIS_protocol/

### 2.4 Data Analysis

#### 2.4.1 NDVI

Following the creation of the orthomosaics the RGB and multispectral images were analyzed via the FieldImageR package in R (Matias, Caraza‐Harter, and Endelman 2020; *R: A Language and Environment for Statistical Computing*, 2023). The package allowed for the NDVI to be extracted using the following equation: (NIR - RED)/(NIR + RED) (Rondeaux, Steven, and Baret 1996).

#### 2.4.2 GDD

To lower error, growing degree days were used to predict the maturity of plants at a given time in the season (Russelle et al. 1984). Through Growing Degree Days (GDD) comparisons, the maturity stage of the corn in 2020 and 2021 was determined in terms of heat units (hours above 10C) or growing degree units (GDU). To compare weather between the two years, GDU were measured using the equation: 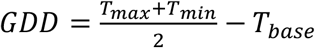 (Baskerville and Emin 1969). T_base_ for corn is 10 degrees Celsius (50 degrees Fahrenheit). This data was used to choose three drone flights and images from each of the two environments that were within approximately 50 GDU of each other (Supplemental Table 1). These three GDU time points labeled by days before and after the flood are: three days pre-flood, six days post-flood, and 26 days-post flood.

**Table 1:** Candidate genes from the GWAS hit on chromosome three along with functional description.

| Chr | Pos (kb) | Locus Name | Description |
| --- | --- | --- | --- |
| 3 | chr3:185762490..185765340 | *Zm00001eb148270* | copper ion binding, hydroquinone: oxygen oxidoreductase activity (laccase8/laccase15) |
| 3 | chr3:185772513..185775910 | *Zm00001eb148280* | copper ion binding, hydroquinone: oxygen oxidoreductase activity (laccase4) |
| 3 | chr3:185836554..185839728 | *Zm00001eb148290* | Scarecrow-like protein (GRAS-transcription factor 23 - gras23) |
| 3 | chr3:185861853..185868788 | *Zm00001eb148300* | Ribonuclease II/R containing protein |
| 3 | chr3:185869509..185870944 | *Zm00001eb148310* | calcium-dependent phospholipid binding (ERP1) |
| 3 | chr3:185906104..185917627 | *Zm00001eb148320* | Transcription factor (myb15) |

#### 2.4.3 GWAS

Genome Wide Association Studies (GWAS) are valuable tools for understanding mechanisms underlying unique phenotypes and identifying markers for breeding. To explore the relationship between maize hybrid genotypes and flood tolerance, GWAS was conducted using the Genomic Association and Prediction Integrated Tool, or GAPIT version 3 with the Mixed Linear and BLINK models (Yu et al., 2006, Huang et al., 2019, Lipka et al., 2012). These models are used to estimate the effect size of each SNP and determine how genotype relates to phenotype. Mixed Linear Models (MLMs) consider both random and fixed effects using the following equation: Y = Si + Q + K + e, where Si is the markers, Q is the population structure, K is the kinship matrix, and e is the residual error. This method accounts for effects from hidden relationships that are not accounted for in naive GWAS. The BLINK model was developed to improve upon FarmCPU by eliminating the assumption that causal genes are evenly distributed across the genome and reducing the number of false positives. Instead of testing against kinship, BLINK tests markers against the associated markers, excluding those in linkage disequilibrium with the most significant marker and repeating that process until none can be excluded. The following equation is used: Y = Si + S + e, where Si is the markers, S is the significant marker, and e is the residual error.

## 3 RESULTS AND DISCUSSION

Initial observation of NDVI raised questions about how values were shifting across the span of the season along with pre- and post-flood. First, a Linear model was fit regressing NDVI on genotype and location in the field (row, range, and row-by-range) to account for distance from the stream and thus flooding duration (supplemental table 2). This showed that both row and range were highly significant (p = 1.19E-05) while the genotype was less important (which was to be expected due to distance from the stream playing a large role in survival). In figure 2, the distribution of NDVI in the flooded field shifted towards lower values of approximately 0.12 during the six days post-flood time. The non-flooded field shows the density shifting to higher NDVI values that reach over 0.6 in a similar maturity point. At the latest time point (26 days post-flood) we see an interesting shift to a bimodal distribution in the flooded field. This first cluster of values centers around 0.09 and the second around 0.19. These initial findings showcased significant variation across the field prompting further investigation.

**Figure 2:**
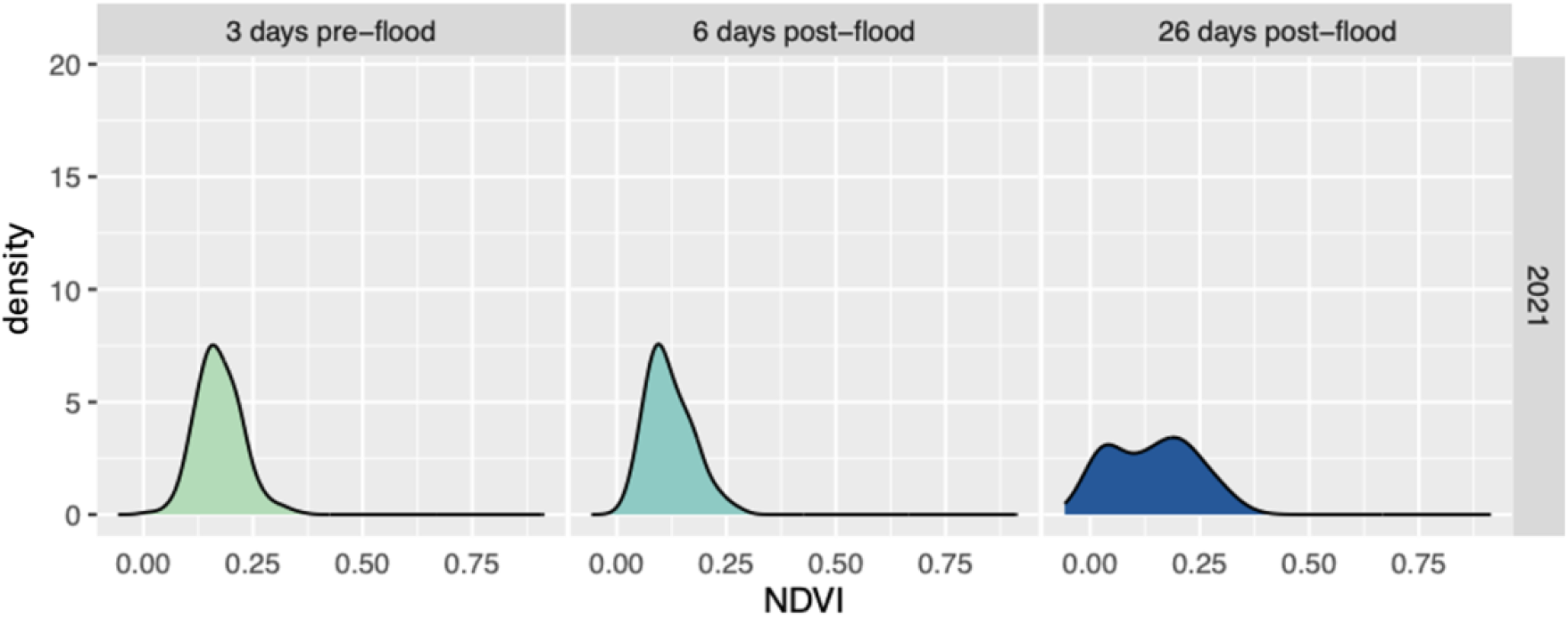
Density plots of the average NDVI in the field. The graphs are separated by year and show the distribution of NDVI values for 3 days pre-flood, 6 days post-flood, and 26 days post-flood.

Before the flooding event, the plants were healthy and growing normally, in fact their NDVI values were higher than those in the non-flooded comparison environment at a similar GDU point in the season (Fig 3. A and D). After flooding, this was reversed with the flooded environment’s NDVI values plummeting while the non-flooded continuing to increase and grow normally (Fig 3. B and E). By the 26 days post-flood time point, stark differences in NDVI values between the flooded and non-flooded environments are even more pronounced (Fig 3. C and F) and the flooded environment shows clear patterns of plants that have recovered to some degree and plants that have not recovered or gotten worse.

**Figure 3:**
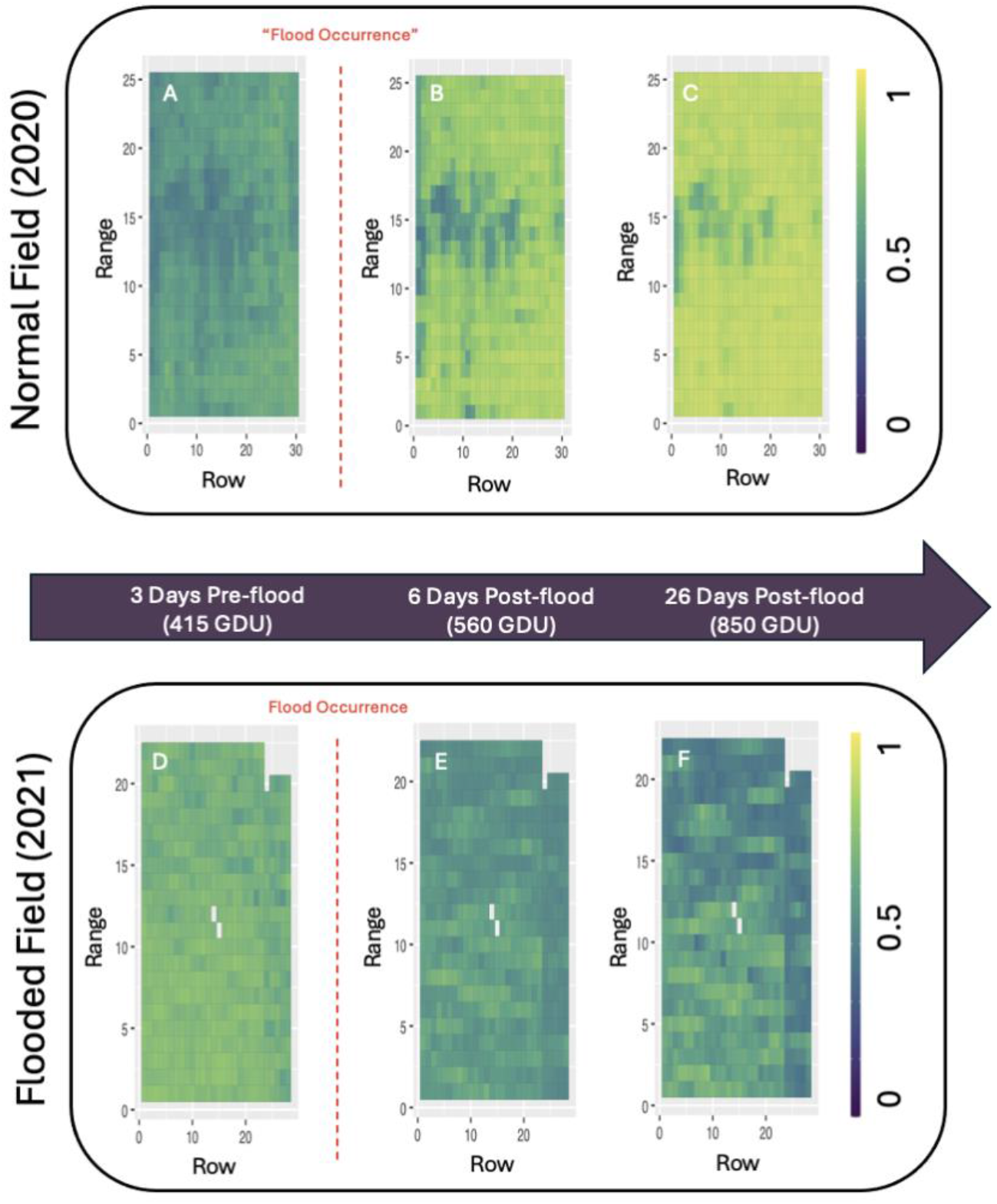
NDVI over the growing season in the flooded and non-flooded control field. Flight dates were chosen via the closest GDU

In Figure 3, stark differences in NDVI values can be seen between 2020 and 2021 in the 26 days post-flood heatmaps (Fig 3 C and F). The flooded field never made a full recovery in comparison to the non-flooded field. The non-flooded field had an average NDVI of 0.862 at 26 days post-flood while the flooded field had a mean value of only 0.535. The NDVI values consistently decreased through every maturity period, reaching their lowest point at 26 days post-flood. The non-flooded field consistently increased in NDVI and maintained values higher than the typical healthy plant threshold of 0.6. The flooded field had 6 plots that increased in NDVI values from 3 days pre-flood to 26 days post-flood, whereas all plots in the control field increased in NDVI value. Understanding the complete damage to the field was not as simple as just looking at the index values. Manual phenotypes such as lodging were taken into account as well.

### Genotypes with Less Lodging were More Successful

After the flooding event occurred, the plants became coated in silty mud and debris (Figure 4B) and damage was observed including the “buggy-whip” phenotype and green snap before the plants reached full maturity (Figure 4C). Drastic decreases in the flooded population’s stand count post-flood were seen (Fig 5C). The pre-flood stand count average was 48.22 plants, while the post-flood stand count average was only 14.96 plants. Overall, there was a 68.98% decrease in standing plants after the flood and only 141 genotypes out of the 410 were able to maintain a stand count above 25 after the flooding event occurred.

**Figure 4:**
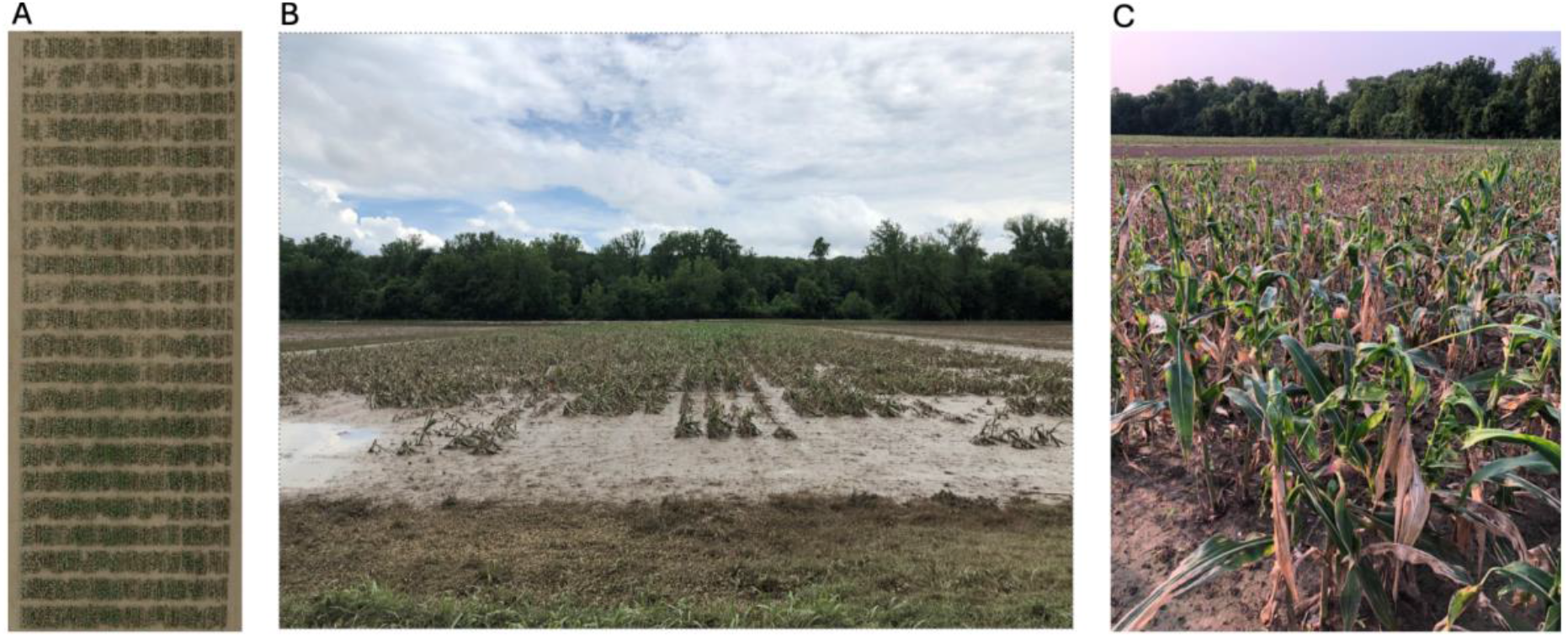
Images of the flooded field the day after the standing water had receded 95% A) gives an aerial image of the field B) Shows the extreme lodging and remaining water/water logging on the field C) “buggy whip” phenotype in the field

**Figure 5:**
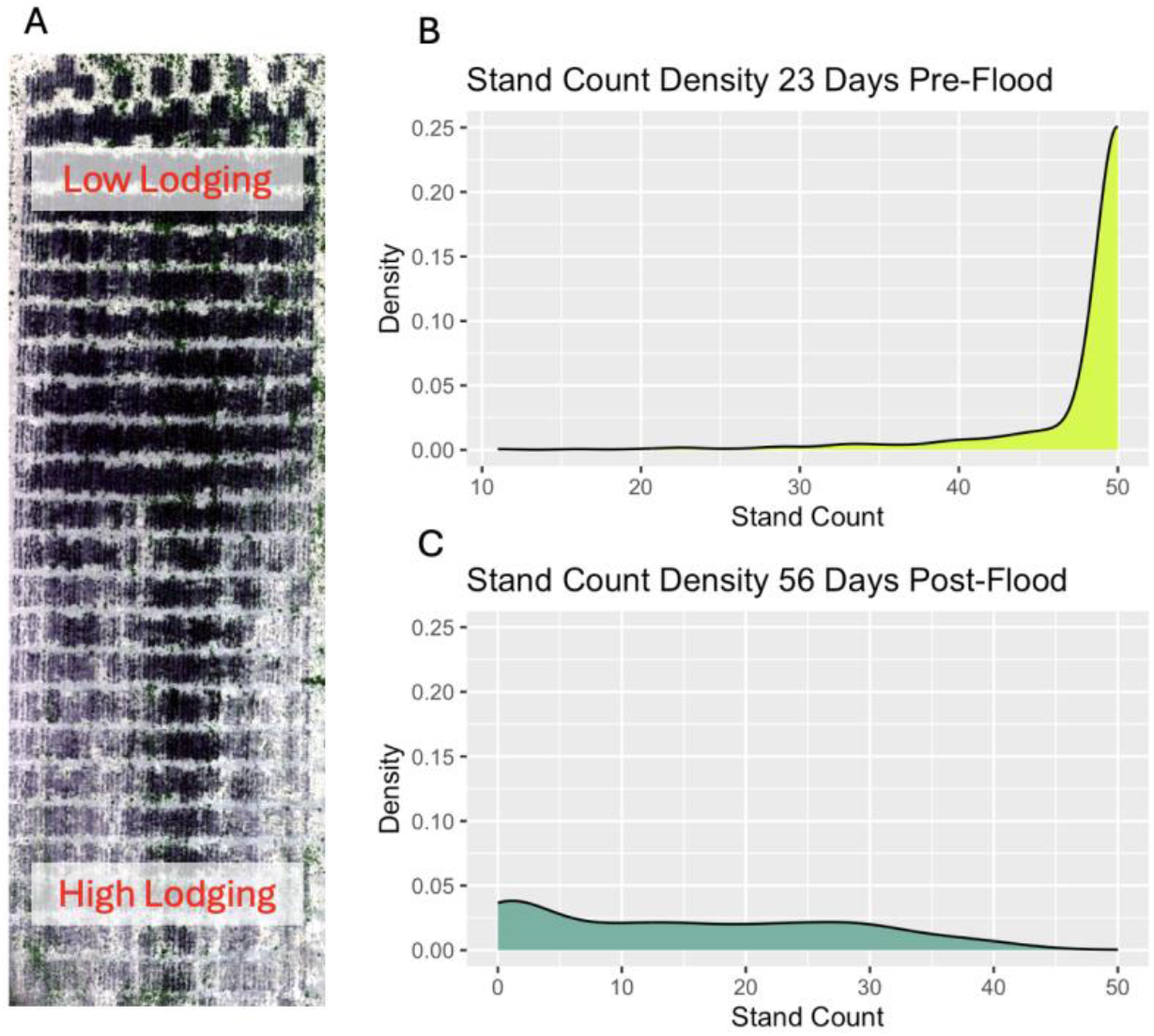
Stand counts of the field before and after the flood. A) Aerial view of the flooded field 6 days-post flood B) 23 Days Pre-Flood stand count distribution (as a density figure) C) 56 Days Post-Flood stand count distribution

Of the six genotypes identified with increasing NDVI values, three had stand counts of 14 plants or higher, and hybrid W10004_1084/PHK76 had a stand count of 32 plants post-flood, more than the post-flood average. The event caused irreversible damage to the plants with limited recovery observed two months post-flooding. This was also witnessed visually across the entirety of the field, with many genotypes exhibiting a correlation between the flooding, lodging, and their extreme decrease in yield. In many sections of the field, grain yield was insufficient for moisture testing and hence precluded yield calculation via a research combine. This in part, is likely due to the misalignment of anthesis and silking of the population that occurred due to the stress of flooding (Supplementary Table 1). The anthesis-silking interval is known to play a large role in the yield of maize (Zaidi et al. 2004).

### Genetic Analysis of the Surviving Lines

The BLINK GWAS results seen in Figure 6 indicate a significant SNP on chromosome 3 from the 26-day post-flood NDVI values from the flooded field. This SNP accounted for 22.11% of the phenotypic variation seen within the flooded field in late-season images. A mixed linear model (MLM) GWAS on the same data shows other SNPs in the vicinity were also within the peak and that the trait had an overall narrow sense heritability of *h*^*2*^ = *0*.*56*. Through the MaizeMiner database, 6 genes of interest were identified using a 200Kbp window around the location of the SNP on chromosome 3. The *Zm00001eb148290* gene is part of the GRAS-domain family of transcription factors and is partially responsible for the developmental regulation of maize (Bai et al. 2022). Specifically, the regulation of Scarecrow functionality. This is responsible for developing C4 anatomy within the plant and endodermis formation in the roots (Slewinski et al. 2012). This leads us to wonder whether this boundary layer’s developmental regulation and formation within the roots could be responsible for elevated NDVI values 26 days post-flood. Along with this, two laccase genes were found, *Zm00001eb148270 and Zm00001eb148280*. These genes are known for their response to stress via monolignin structure adaptation (Wang et al. 2024). Lignin in maize is known to increase with maturity (He et al. 2018). Furthermore, *Zm00001eb148310* is annotated as “elicitor-responsive protein 1” by the National Center for Biotechnology Information (NCBI) thought to elicit a drought stress responses in maize (Lin et al. 2024). A deeper investigation into the functionality of these genes under flooding stress needs to be conducted to fully identify what allowed plants to maintain high NDVI levels post-flood.

**Figure 6:**
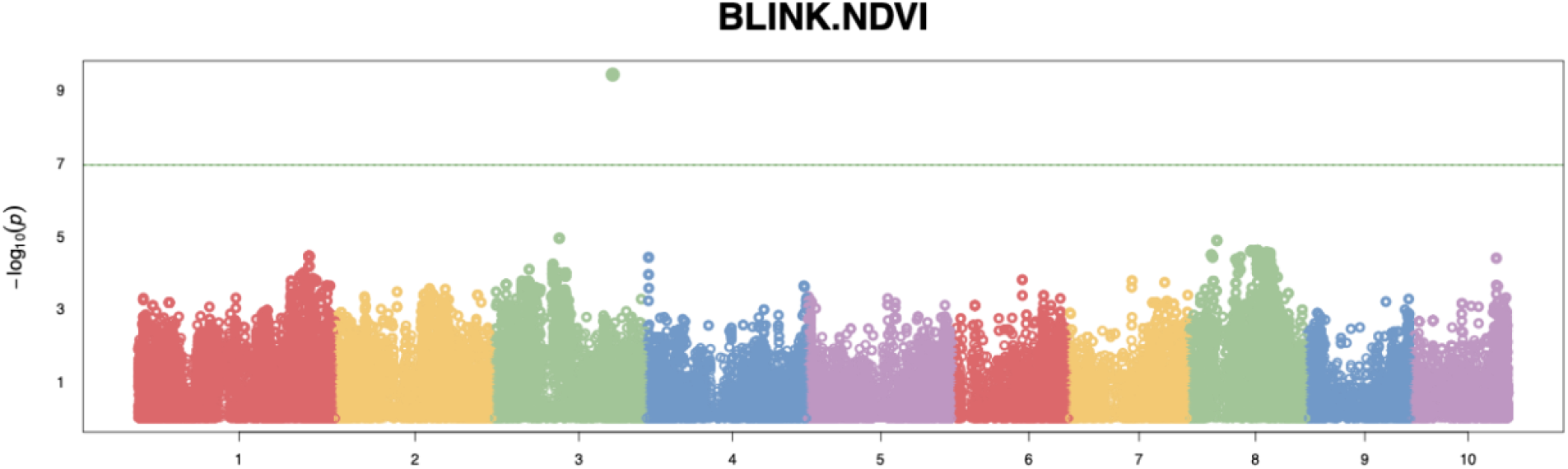
Manhattan plot of the NDVI at 26 days post-flood created via GAPIT’s BLINK model

## Conclusions

Overall, the utility of drones in capturing relevant flood data was clearly demonstrated. This approach allows researchers and farmers to analyze damage without the risk of harm that large-scale flooding can impose on humans. It also benefits the discovery and identification of potential genes of interest for breeders. Drones allow phenotyping to take place in an aerial, shoot-based setting instead of underground or in a simulated environment. However, some reservations should still be held about the external factors that can influence the data shown. While temperature and plant maturity were considered during cross-season analysis, factors such as management, soil characteristics, and pathogens were not measured, making it premature to draw explicit conclusions from the flooded and non-flooded field comparison.

With this in mind, we have shown that the use of UAVs can accurately collect data to be used in the extraction of multispectral indices to quantify plant physiological responses to flooding stress. This methodology highlighted areas of the field that were photosynthetically active and indicated the overall health of each plot in our experiment. Further NDVI gave us a significant hit in our GWAS analysis that was responsible for over one-fifth of the phenotypic variance that was seen amongst plants and 6 potential genes of interest. In all, this gives tangible value to the utilization of drones to assess the damage and recovery of flooded maize.

## Abbreviations

G2F: Genomes to Fields
GxE: Genome by Environment
GDD: Growing Degree Days
GDU: Growing Degree Units
GRAS: Gibberellic Acid Insensitive, Repressor of GA1 and SCARECROW
GWAS: Genome Wide Association Study
MLM: Mixed Linear Model
MODIS: Moderate Resolution Imaging Spectroradiometer
NCBI: National Center for Biotechnology Information
NDVI: Normalized Difference Vegetative Index
NIR: Near Infrared
RGB: Red Green Blue
SNP: Single Nucleotide Polymorphism
UAV: Unoccupied Aerial Vehicle

## ACKNOWLEDGMENTS

This work was funded by the United States Department of Agriculture – Agricultural Research Service.

## DATA AVAILABILITY

All data and code utilized in the analysis is available in the following repository: https://github.com/mnmzbd24/Flood_Tolerance

## CONFLICT OF INTEREST

The authors declare no conflict of interest

## Notes

https://www.protocols.io/edit/qgis-gridding-protocol-dc872zzn.

https://github.com/mnmzbd24/Flood_Tolerance

### Competing Interest Statement

The authors have declared no competing interest.

## REFERENCES

Bai, Yibo, Hui Liu, Kaikai Zhu, and Zong-Ming Cheng. 2022. “Evolution and Functional Analysis of the GRAS Family Genes in Six Rosaceae Species.” BMC Plant Biology 22(1):569. doi: 10.1186/s12870-022-03925-x.

Baskerville, G. L., and P. Emin. 1969. “Rapid Estimation of Heat Accumulation from Maximum and Minimum Temperatures.” Ecology 50(3):514–17. doi: 10.2307/1933912.

Genomes to Fields. 2024. “GxE Field Experiment 2024 SOP.”

He, Yuan, Thibaut Mb Mouthier, Mirjam A. Kabel, Jan Dijkstra, Wouter H. Hendriks, Paul C. Struik, and John W. Cone. 2018. “Lignin Composition Is More Important than Content for Maize Stem Cell Wall Degradation.” Journal of the Science of Food and Agriculture 98(1):384–90. doi: 10.1002/jsfa.8630.

Huang, M., Liu, X., Zhou, Y., Summers, R. M., & Zhang, Z. (2019). BLINK: A package for the next level of genome-wide association studies with both individuals and markers in the millions. GigaScience, 8(2), giy154. 10.1093/gigascience/giy154

Jiménez, Juan De La Cruz, Angelika Mustroph, Ole Pedersen, Daan A. Weits, and Romy Schmidt-Schippers. 2024. “Flooding Stress and Responses to Hypoxia in Plants” edited by S. Shabala. Functional Plant Biology 51(4). doi: 10.1071/FP24061.

Kaur, Gurpreet, Brendan Zurweller, Peter Motavalli, and Kelly Nelson. 2019. “Screening Corn Hybrids for Soil Waterlogging Tolerance at an Early Growth Stage.” Agriculture 9(2):33. doi: 10.3390/agriculture9020033.

Lin, Yi-Hsuan, Ya-Ning Zhou, Xiao-Gui Liang, Yu-Ka Jin, Zu-Dong Xiao, Ying-Jun Zhang, Cheng Huang, Bo Hong, Zhen-Yuan Chen, Shun-Li Zhou, and Si Shen. 2024. “Exogenous Methylglyoxal Alleviates Drought-Induced ‘Plant Diabetes’ and Leaf Senescence in Maize” edited by C. Foyer. Journal of Experimental Botany 75(7):1982–96. doi: 10.1093/jxb/erad503.

Lindsey, Alexander J., Paul R. Carter, and Peter R. Thomison. 2021. “Impact of Imposed Root Lodging on Corn Growth and Yield.” Agronomy Journal 113(6):5054–62. doi: 10.1002/agj2.20848.

Lipka, A. E., Tian, F., Wang, Q., Peiffer, J., Li, M., Bradbury, P. J., Gore, M. A., Buckler, E. S., & Zhang, Z. (2012). GAPIT: Genome association and prediction integrated tool. Bioinformatics, 28(18), 2397–2399. 10.1093/bioinformatics/bts444

Liu, Yuan, Chenwei Nie, Zhen Zhang, ZiXu Wang, Bo Ming, Jun Xue, Hongye Yang, Honggen Xu, Lin Meng, Ningbo Cui, Wenbin Wu, and Xiuliang Jin. 2023. “Evaluating How Lodging Affects Maize Yield Estimation Based on UAV Observations.” Frontiers in Plant Science 13:979103. doi: 10.3389/fpls.2022.979103.

Mano, Y., F. Omori, T. Takamizo, B. Kindiger, R. McK. Bird, and C. H. Loaisiga. 2006. “Variation for Root Aerenchyma Formation in Flooded and Non-Flooded Maize and Teosinte Seedlings.” Plant and Soil 281(1–2):269–79. doi: 10.1007/s11104-005-4268-y.

Matias, Filipe Inácio, Maria V. Caraza‐Harter, and Jeffrey B. Endelman. 2020. “FIELDimageR: An R Package to Analyze Orthomosaic Images from Agricultural Field Trials.” The Plant Phenome Journal 3(1):e20005. doi: 10.1002/ppj2.20005.

MicaSense RedEdge MX processing workflow (including reflectance calibration) in Agisoft metashape professional. Agisoft Helpdesk Portal. (2024, November 12). https://agisoft.freshdesk.com/support/solutions/articles/31000148780-micasense-rededge-mx-processing-workflow-including-reflectance-calibration-in-agisoft-metashape-pro#Build-Dense-Cloud

QGIS.org, %%Y. QGIS 3.34. Geographic Information System User Guide. QGIS Association. Electronic document: https://docs.qgis.org/3.34/en/docs/user_manual/index.html

R: A language and environment for statistical computing. (2023). [Computer software]. R Foundation for Statistical Computing. https://www.R-project.org/

Razzaq, Ali, Shabir Hussain Wani, Fozia Saleem, Min Yu, Meixue Zhou, and Sergey Shabala. 2021. “Rewilding Crops for Climate Resilience: Economic Analysis and de Novo Domestication Strategies” edited by C. Foyer. Journal of Experimental Botany 72(18):6123–39. doi: 10.1093/jxb/erab276.

Ren, Baizhao, Jiwang Zhang, Shuting Dong, Peng Liu, and Bin Zhao. 2016. “Root and Shoot Responses of Summer Maize to Waterlogging at Different Stages.” Agronomy Journal 108(3):1060–69. doi: 10.2134/agronj2015.0547.

Rondeaux, Geneviève, Michael Steven, and Frédéric Baret. 1996. “Optimization of Soil-Adjusted Vegetation Indices.” Remote Sensing of Environment 55(2):95–107. doi: 10.1016/0034-4257(95)00186-7.

Rosenzweig, Cynthia, Francesco N. Tubiello, Richard Goldberg, Evan Mills, and Janine Bloomfield. 2002. “Increased Crop Damage in the US from Excess Precipitation under Climate Change.” Global Environmental Change 12(3):197–202. doi: 10.1016/S0959-3780(02)00008-0.

Russelle, M. P., W. W. Wilhelm, R. A. Olson, and J. F. Power. 1984. “Growth Analysis Based on Degree Days ^1^.” Crop Science 24(1):28–32. doi: 10.2135/cropsci1984.0011183×002400010007x.

Shrestha, Ranjay, Liping Di, Genong Yu, Yuanzheng Shao, Lingjung Kang, and Bei Zhang. 2013. “Detection of Flood and Its Impact on Crops Using NDVI - Corn Case.” Pp. 200– 204 in 2013 Second International Conference on Agro-Geoinformatics (Agro-Geoinformatics). Fairfax, VA, USA: IEEE.

Slewinski, Thomas L., Alyssa A. Anderson, Cankui Zhang, and Robert Turgeon. 2012. “Scarecrow Plays a Role in Establishing Kranz Anatomy in Maize Leaves.” Plant and Cell Physiology 53(12):2030–37. doi: 10.1093/pcp/pcs147.

Southworth, Jane, J. C. Randolph, M. Habeck, O. C. Doering, R. A. Pfeifer, D. G. Rao, and J. J. Johnston. 2000. “Consequences of Future Climate Change and Changing Climate Variability on Maize Yields in the Midwestern United States.” Agriculture, Ecosystems & Environment 82(1–3):139–58. doi: 10.1016/S0167-8809(00)00223-1.

Sweet, Dorothy D., Sara B. Tirado, Nathan M. Springer, Candice N. Hirsch, and Cory D. Hirsch. 2022. “Opportunities and Challenges in Phenotyping Row Crops Using Drone‐based RGB Imaging.” The Plant Phenome Journal 5(1):e20044. doi: 10.1002/ppj2.20044.

Tian, Li-xin, Wen-shuang Bi, Xiao-song Ren, Wen-long Li, Lei Sun, and Jing Li. 2020. “Flooding Has More Adverse Effects on the Stem Structure and Yield of Spring Maize (Zea Mays L.) than Waterlogging in Northeast China.” European Journal of Agronomy 117:126054. doi: 10.1016/j.eja.2020.126054.

Tirado, Sara B., Candice N. Hirsch, and Nathan M. Springer. 2021. “Utilizing Temporal Measurements from UAVs to Assess Root Lodging in Maize and Its Impact on Productivity.” Field Crops Research 262:108014. doi: 10.1016/j.fcr.2020.108014.

United States Department of Agriculture - Natural Resources Conservation Service. n.d. “Web Soil Survey.”

United States Department of Agriculture - Risk Management Agency. n.d. “Cause of Loss Historical Data Files.”

Victor McDaniel, R. Wayne Skaggs, and Lamyaa M. Negm. 2016. “Injury and Recovery of Maize Roots Affected by Flooding.” Applied Engineering in Agriculture 32(5):627–38. doi: 10.13031/aea.32.11633.

Wang, Tonghan, Yang Liu, Kunliang Zou, Minhui Guan, Yutong Wu, Ying Hu, Haibing Yu, Junli Du, and Degong Wu. 2024. “The Analysis, Description, and Examination of the Maize LAC Gene Family’s Reaction to Abiotic and Biotic Stress.” Genes 15(6):749. doi: 10.3390/genes15060749.

Yu, J., Pressoir, G., Briggs, W. H., Vroh Bi, I., Yamasaki, M., Doebley, J. F., McMullen, M. D., Gaut, B. S., Nielsen, D. M., Holland, J. B., Kresovich, S., & Buckler, E. S. (2006). A unified mixed-model method for association mapping that accounts for multiple levels of relatedness. Nature Genetics, 38(2), 203–208. 10.1038/ng1702

Zaidi, Pervez H., S. Rafique, P. K. Rai, N. N. Singh, and G. Srinivasan. 2004. “Tolerance to Excess Moisture in Maize (Zea Mays L.): Susceptible Crop Stages and Identification of Tolerant Genotypes.” Field Crops Research 90(2–3):189–202. doi: 10.1016/j.fcr.2004.03.002.

