## SUPPLEMENTARY MATERIAL for "Drone-Based Identification of Flood-Tolerant Maize via Multispectral Imaging: A Real-World Case Study"

| Timeframe | Observation |
| --- | --- |
| 3 days pre-flood | Healthy V5 maize growing uniformly across the field |
| 1 day post-flood | Covered in silt causing the plants to look as thought they were painted grey |
| 14 days post-flood | Majority of the field entered the reproductive phase. Many plants went into anthesis and shortly after silking. But the lack of synchronization revealed yield problems later in the season |
| 14 days post-flood | Cups of mud remained in the whorls forcing stems to snap as elongation continued |
| 14 days post-flood | Many tassels became stuck in their sheaths causing a "buggy-whip" phenotype to occur |

**Supplementary Table 1:** Visual observations made in the field throughout the growing season denoted via a pre- and post-flood timeline

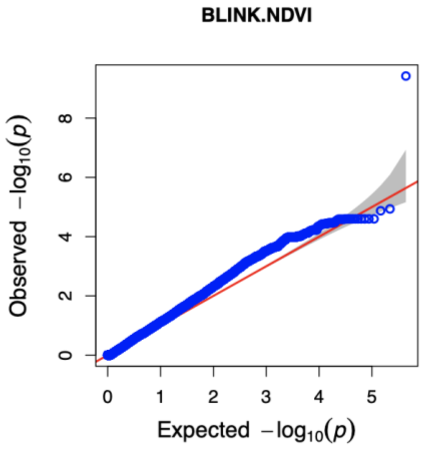


**Supplementary Figure 1:** QQ Plot from BLINK GWAS analysis

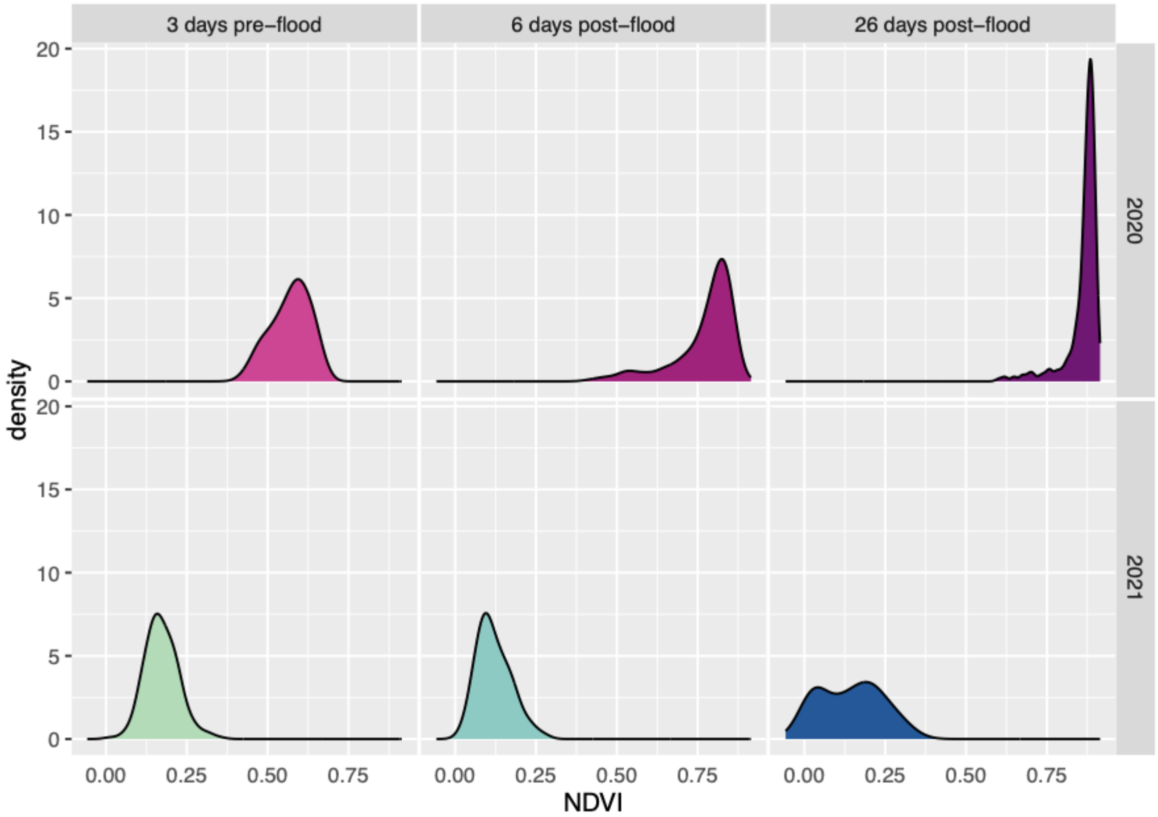


**Supplementary Figure 2:** Density plots of the flooded (2021) and non-flooded (2020) fields.

| Field Treatment | 3 days pre-flood | 6 days post-flood | 26 days post-flood |
|---|---|---|---|
| Normal | 387.82 | 549.53 | 855.63 |
| Flood | 444.72 | 573.62 | 848.74 |
| Difference | -56.9 | -24.09 | 6.89 |

**Supplementary Table 2:** Exact GDU values for the flights chosen for comparison and their differences

| Term | sumsq | df | statistic | p.value |
|---|---|---|---|---|
| (Intercept) | 1.292737402 | 1 | 224.2440253 | 2.15E-33 |
| Pedigree | 2.416138602 | 420 | 0.997891033 | 0.513422151 |
| Exp_row | 0.247763544 | 1 | 42.97817504 | 5.72E-10 |
| Exp_range | 0.33569211 | 1 | 58.23065842 | 1.33E-12 |
| Exp_row:Exp_range | 0.117061513 | 1 | 20.30601492 | 1.19E-05 |
| Residuals | 1.031911529 | 179 | NA | NA |

**Supplementary Figure 1:** ANOVA table for linear model of the row, range, and pedigree of the plants in comparison to their final NDVI values.
